# Inactivation of SARS-CoV-2 and influenza A virus by spraying hypochlorous acid solution and hydrogen peroxide solution in the form of Dry Fog

**DOI:** 10.1101/2021.12.13.472413

**Authors:** Masahiro Urushidani, Akira Kawayoshi, Tomohiro Kotaki, Keiichi Saeki, Yasuko Mori, Masanori Kameoka

## Abstract

Severe acute respiratory syndrome coronavirus 2 (SARS-CoV-2), the causative agent of coronavirus disease 2019 (COVID-19), is transmitted by droplet and contact infection. SARS-CoV-2 that adheres to environmental surfaces remains infectious for several days. We herein attempted to inactivate SARS-CoV-2 and influenza A virus adhering to an environmental surface by spraying aerosolized hypochlorous acid solution and hydrogen peroxide solution in the form of Dry Fog (fog that does not wet objects even if touched). SARS-CoV-2 and influenza virus were dried on plastic plates and placed into a test chamber for inactivation by the Dry Fog spraying of disinfectants. The results obtained showed that Dry Fog spraying inactivated SARS-CoV-2 and influenza A virus in time- and exposed disinfectant amount-dependent manners. SARS-CoV-2 was more resistant to the virucidal effects of aerosolized hypochlorous acid solution and hydrogen peroxide solution than influenza A virus; therefore, higher concentrations of spray solutions were required to inactivate SARS-CoV-2 than influenza A virus. The present results provide important information for the development of a strategy that inactivates SARS-CoV-2 and influenza A virus on environmental surfaces by spatial spraying.

## Introduction

Coronavirus disease 2019 (COVID-19) continues to spread worldwide, with more than 266 million individuals being infected and more than 5.26 million dying to date [1]. The World Health Organization (WHO) has recommended a number of countermeasures to the public, including getting vaccinated, avoiding 3Cs (spaces that are closed, crowded, or involve close contact), wearing a properly fitting mask when physical distancing is not possible and in poorly ventilated settings, and frequently cleaning hands with an alcohol-based hand rub or soap and water [2]; however, the pandemic continues.

Severe acute respiratory syndrome coronavirus 2 (SARS-CoV-2) is the causative agent of COVID-19. Viral transmission is established by inhaling droplets or aerosols containing the virus excreted from infected individuals in a 3Cs setting or by touching the eyes, nose, or mouth with hands contaminated with the virus adhering to environmental surfaces [3]. Previous studies reported the high stability of SARS-CoV-2 adhering to environmental surfaces [4, 5], and its stability was shown to be higher than those of SARS-CoV and influenza A virus [6, 7]. Therefore, the disinfection of environmental surfaces is indispensable as an infection control measure. However, it is not realistic to frequently and manually disinfect the surfaces of large spaces, such as a train station or airport, because of the manpower and time required. Although the spraying of a disinfectant is an alternative spatial disinfection method, it is not recommended by WHO due to its effects on the human body [8]. Furthermore, the Center for Disease Control and Prevention of the United States of America (USA) does not recommend the spraying of disinfectants in hospital rooms because due to the lack of scientific verification on effective, reliable, and safe spatial spraying methods for disinfectants, it is not regarded as an adequate method for decontaminating air and surfaces [9].

Therefore, the present study examined the effectiveness of inactivating SARS-CoV-2 virus adhering to plastic microplates by spraying disinfectant in the form of Dry Fog, which is defined as an aerosol with a Sauter mean droplet diameter ≤10 μm and maximum droplet diameter ≤50 μm. Its virucidal effects on influenza A virus, which is a common envelope virus transmitted through droplets and contact transmission worldwide, were also investigated. In consideration of the effects of residues after spraying on the human body, we tested hypochlorous acid solution and hydrogen peroxide solution, which leave almost no residue on environmental surfaces after spraying. Hypochlorous acid solution is an aqueous solution that contains hypochlorous acid (HOCl) as the main component. Hypochlorous acid solution may be prepared by dissolving sodium hypochlorite in water with adjustments to a weak acidic pH. The main component of the weakly acidic solution is HOCl, while that of an alkaline solution is hypochlorite ions (OCl-). HOCl exerts stronger bactericidal effects than OCl- [10]. Therefore, a weakly acidic (pH 6.5) hypochlorous acid solution was sprayed in the form of Dry Fog in the present study.

## Materials and methods

### Preparation of spray disinfectants

Hypochlorous acid solution and hydrogen peroxide solution were prepared as disinfectants to be sprayed in the form of Dry Fog for the inactivation of viruses. Commercially available, weakly acidic (pH 6.5) hypochlorous acid solution with a free available chlorine (FAC) concentration (the total of HOCl and OCl-concentrations) [10] of 250 ppm (Super Jiasui; HSP Corporation, Okayama, Japan) and a solution diluted by distilled water with a FAC concentration of 125 ppm were used. To prepare a solution with a FAC concentration of 8,700 ppm, sodium hypochlorite (Hayashi Pure Chemical Ind., Ltd., Osaka, Japan) was dissolved in distilled water, followed by an adjustment of the pH of the solution to 6.5 with HCl (Hayashi Pure Chemical Ind., Ltd.). Commercially available hydrogen peroxide solution (56,400 ppm; Part 1, Decon7; Decon7 Systems LLC., Texas, USA) and solutions diluted by distilled water with hydrogen peroxide concentrations of 11,280, 5,640, 2,820, and 1,410 ppm were prepared. Regarding the negative control, distilled water was sprayed in the form of Dry Fog.

### Cells, viruses, and preparation of dried virus samples

VeroE6/TMPRESS2 cells (JCRB1819) [11] were maintained in Dulbecco’s modified Eagle medium (DMEM) (Nacalai Tesque, Inc., Kyoto, Japan) supplemented with 10% fetal bovine serum (Sigma-Aldrich, Merck, Kenilworth, New Jersey, USA) and 1 mg/ml of G418 (Sigma-Aldrich) (complete DMEM), while MDCK cells were maintained in Eagle’s minimum essential medium (MEM) (MEM1, Nissui Pharmaceutical Co., Ltd., Tokyo, Japan) supplemented with 10% FBS in a CO_2_ incubator. The SARS-CoV-2 Wuhan strain, SARS-CoV-2/Hu/DP/Kng/19-020 was propagated by infecting VeroE6/TMPRESS2 cells and cultured in FBS-free DMEM supplemented with 1 mg/ml of G418 for 24 hours. The influenza A H1N1 strain, A/Puerto Rico/8/1934 was propagated by infecting MDCK cells and cultured in FBS-free MEM supplemented with 2 μg/ml of acetylated trypsin (Sigma-Aldrich) for 3 days. Viral supernatants were clarified by centrifugation, aliquoted, and stored at −85 °C. Ten-fold serially diluted viral supernatants were incubated with VeroE6/TMPRESS2 or MDCK cells for 3 or 4 days, respectively, for viral titration, and median tissue culture infectious doses (TCID_50_) were measured following the fixation of cells with 5% formaldehyde in phosphate-buffered saline (PBS) and staining with 0.5% crystal violet in 20% EtOH. TCID_50_ values below the detection limit (3.16 × 10^2^ TCID_50_/ml or 2.5 log10 TCID_50_/ml) were assigned to half of the detection limit, equivalent to 1.58 × 10^2^ TCID_50_/ml or 2.2 log10 TCID_50_/ml, because substituting the value to half of the detection limit was previously shown to be less biased than substitution to zero or the detection limit [12]. SARS-CoV-2 (1.2 × 10^5^ TCID_50_ in 5 μl) and influenza A virus (2.8 × 10^6^ TCID_50_ in 5 μl) were added to 96-well flat-bottomed microplates (Corning Japan, Shizuoka, Japan), air-dried for 10-15 minutes using a small electric fan, and subjected to inactivation experiments. In addition, for some experiments, 5 μl of artificial saliva (Saliveht aerosol; Teijin Pharma, Tokyo, Japan) or PBS was mixed with 5 μl of viral supernatants prior to air-drying samples.

### Preparation of the test chamber for spraying

A closed test chamber was prepared to fill the space with Dry Fog. The detailed settings of the chamber are shown in Figure 1. The chamber size was 500 × 700 × 300 mm (height × width × depth) and made of acrylic, and it was set in a biosafety cabinet. A sliding door was set on the front of the chamber for the handling of samples inside the chamber. A sprayer equipped with an impinging-jet atomizing nozzle [13] [AE-1 (03C), AKIMist®”E”; H. Ikeuchi & Co., Ltd., Osaka, Japan] was used to spray a disinfectant into the space. To generate aerosolized disinfectants in the form of Dry Fog, 0.3 MPa compressed air was supplied to the sprayer from a compressor (0.2LE-8SB0; Hitachi Industrial Equipment Systems Co., Ltd., Tokyo, Japan). The spray capacity was 2.3 liters per hour and the Sauter mean droplet diameter was 7.5 μm. In the virus inactivation experiment using Dry Fog spraying, four 90-mm Petri dishes containing 20 ml of distilled water, a thermo-hygrometer, and a 96-well microplate containing dried viral samples were placed in the chamber (Fig. 1).

**Fig. 1.**
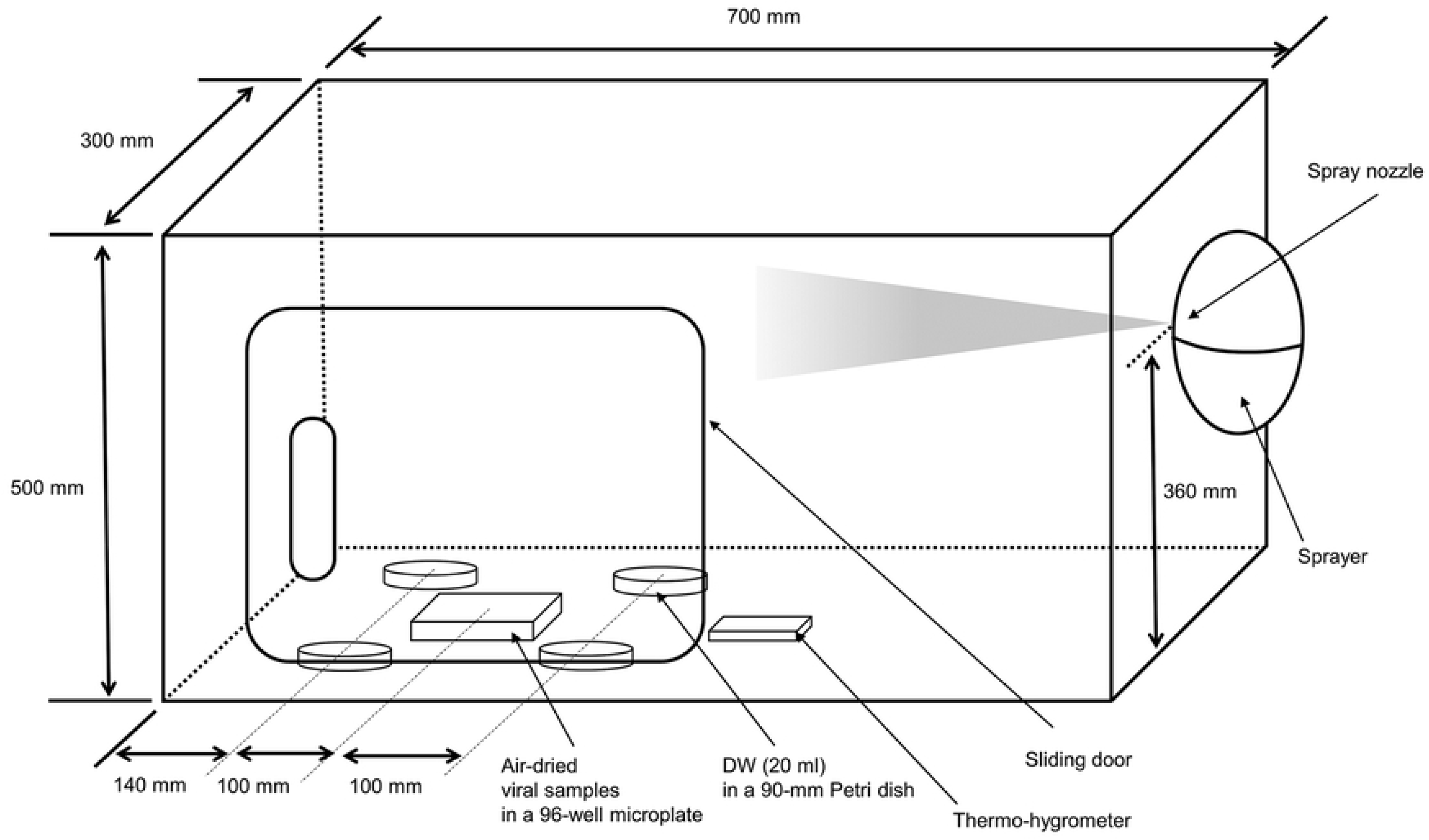
Schematic illustration of the test chamber. The chamber was made of acrylic at the indicated size, and a sliding door was set on the front. A sprayer that generates the Dry Fog of spray solution was set on the right side of the chamber at a height of 360 mm. In virus inactivation experiments, four 90-mm Petri dishes containing 20 ml of distilled water (DW), a thermo-hygrometer, and a 96-well microplate containing air-dried viral samples were placed in the chamber, as indicated.

### Virus inactivation experiment by Dry Fog spraying and measurement of the exposed disinfectant amount to samples

The times at which spatial spraying, the termination of virus inactivation operation, and the measurement of the exposed disinfectant amount to the environmental surface were performed are indicated in Figure 2. Spatial spraying was conducted as follows. A disinfectant was sprayed for 5 seconds at the start of the experiment and left to stand for 4 minutes. Spraying was then repeated 3 more times for 2.5 seconds each, and left to stand for 4 minutes after each spraying. Spraying was performed 4 times, namely, 0, 4, 8, and 12 minutes after the initiation of the experiment, and the total experimental period was 16 minutes. This spraying operation allowed for the continuous generation of Dry Fog without unnecessarily moistening the environmental surface in the chamber, and Dry Fog did not fade. The virus inactivation reaction by spraying a disinfectant was terminated by resuspending air-dried viral samples in 200 μl of DMEM containing a neutralizer of the disinfectant being tested. DMEM containing 0.1 M sodium thiosulfate (Hayashi Pure Chemical Ind., Ltd.) was used as a neutralizer for hypochlorous acid solution, while DMEM containing 0.1 mg/ml of catalase (Nacalai Tesque, Inc.) was used for hydrogen peroxide solution. The residual infectious titer of viral samples after the inactivation experiment was evaluated by measuring the TCID_50_ value, as described above. The exposed disinfectant amount of the spray to viral samples was assessed by collecting the droplet-shaped disinfectant that had fallen into distilled water in the 90-mm Petri dish placed in the chamber, and measuring the disinfectant concentration dissolved in distilled water. The FAC concentration when hypochlorous acid solution was sprayed was measured by DPD absorptiometry using a residual chlorine meter (HI96771; Hanna Instruments, Chiba, Japan). The hydrogen peroxide concentration when hydrogen peroxide solution was sprayed was measured by 4-aminoantipyrine absorptiometry using an enzyme with a hydrogen peroxide concentration meter (DPM2-H_2_O_2_; Kyoritsu Chemical-Check Lab., Corp., Kanagawa, Japan). Concentrations in appropriately diluted samples with distilled water were within the detection ranges of the measuring instruments and reagents. In addition, air temperature and humidity in the chamber were monitored during experiments.

**Fig. 2.**
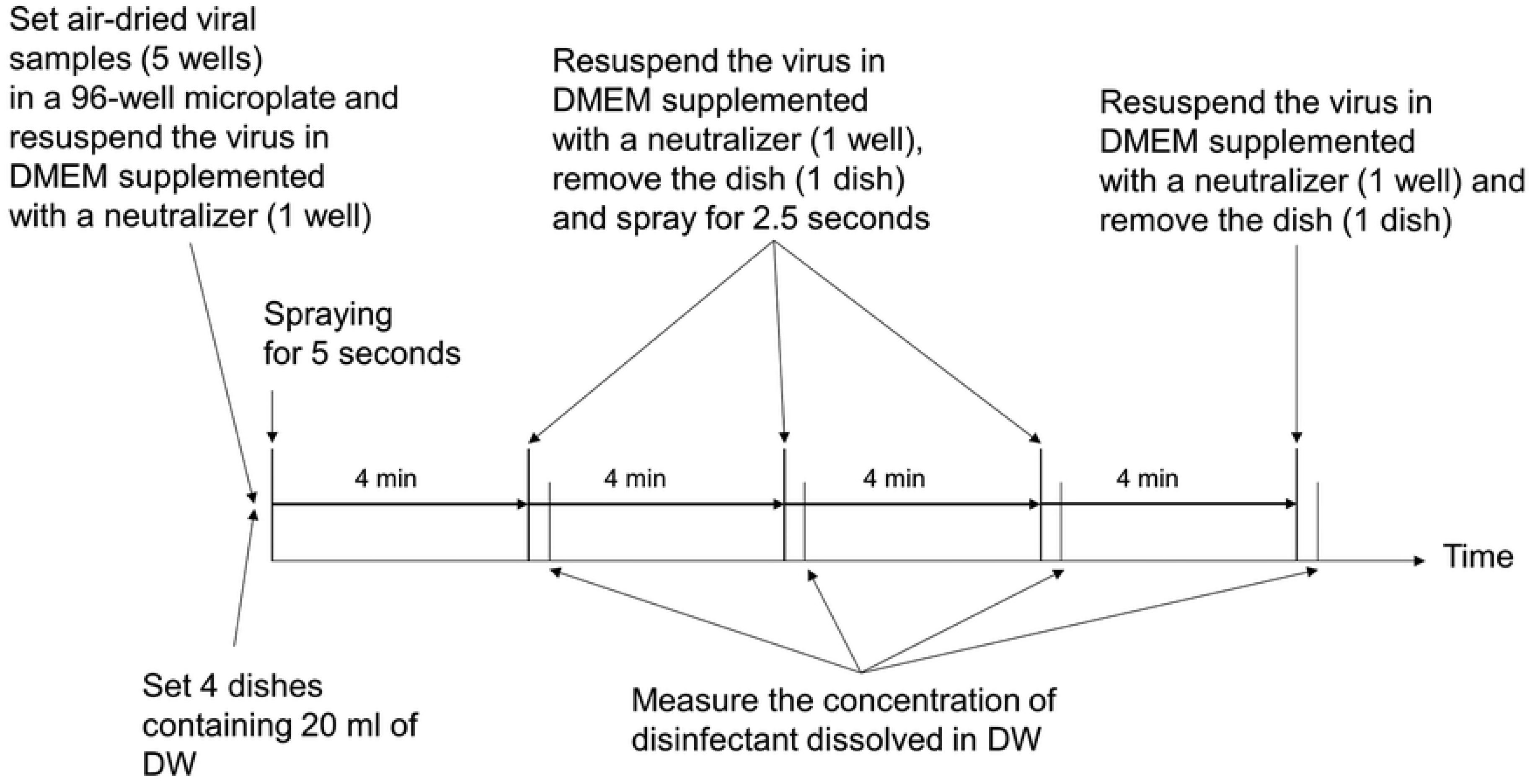
Flow diagram of the virus inactivation experiment. The times at which spatial spraying was performed, the virus inactivation operation was terminated, and the exposed disinfectant amount to the environmental surface was measured are as indicated. A detailed procedure is described in the Materials and methods.

Experiments were performed as follows. After the spraying of disinfectant and leaving it to stand for 4 minutes, the sliding door of the chamber was opened, viral samples were resuspended with DMEM containing a neutralizer, the Petri dish containing distilled water was removed from the chamber, the sliding door was closed, and the disinfectant was sprayed and left to stand again for 4 minutes. The concentration of the disinfectant in distilled water in the Petri dish removed from the chamber was then measured. The virus inactivation reaction was terminated by resuspending samples in DMEM containing a neutralizer before spraying three more times. Therefore, the exposed disinfectant amount to viral samples was measured 5 times in each set of experiments, i.e., at 0 minutes, 4 minutes (5 seconds of spraying), 8 minutes (7.5 seconds of spraying), 12 minutes (10 seconds of spraying), and 16 minutes (12.5 seconds of spraying).

### Statistical analysis

Statistical analyses were performed using the standard function of GraphPad Prism 8 software (GraphPad Software, San Diego, California) with a 2-way ANOVA, unpaired *t*-test, a non-linear regression (curve fit) analysis, or simple linear regression analysis.

## Results

### The Dry fog spraying of hypochlorous acid solution and hydrogen peroxide solution inactivated SARS-CoV-2 and influenza A virus in a time-dependent manner

We examined changes in the viral infectious titer upon spraying Dry Fog with various concentrations of hypochlorous acid solution (FAC concentrations of 8,700, 250, and 125 ppm), hydrogen peroxide solution (56,400, 11,280, 5640, 2820, and 1410 ppm of hydrogen peroxide), or distilled water. Spraying experiments were initially conducted using a commercially available hypochlorous acid solution (250 ppm; Super Jiasui; HSP); however, SARS-CoV-2 was not inactivated (Fig. 3A). A previous study reported that when viral culture fluid was mixed with 35 ppm hypochlorous acid solution, SARS-CoV-2 was effectively inactivated [14]; therefore, we calculated the FAC concentration needed to inactivate an air-dried virus settled in the wells (0.32 cm^2^) of 96-well microplates. The result obtained revealed that 8,700 ppm hypochlorous acid solution sprayed as Dry Fog was required to inactivate SARS-CoV-2 under our experimental conditions. Figure 3 shows the time course for the virus infectious titer under experimental conditions that inactivated the virus. The viral titer of SARS-CoV-2 was not reduced by 250 ppm hypochlorous acid solution (Fig. 3A), whereas that of influenza A virus was effectively decreased in a time-dependent manner (Fig. 3B). Furthermore, 8,700 ppm hypochlorous acid solution effectively reduced the infectious titer of SARS-CoV-2 (Fig. 3A). Moreover, 56,400 ppm hydrogen peroxide solution reduced the infectious titer of SARS-CoV-2 (Fig. 3C), while 11,280 ppm hydrogen peroxide solution decreased the viral titer of influenza A virus (Fig. 3D). The Dry Fog spraying of distilled water did not reduce the viral infectivity of SARS-CoV-2 or influenza A virus regardless of the elapsed time (Fig. 3).

**Fig. 3.**
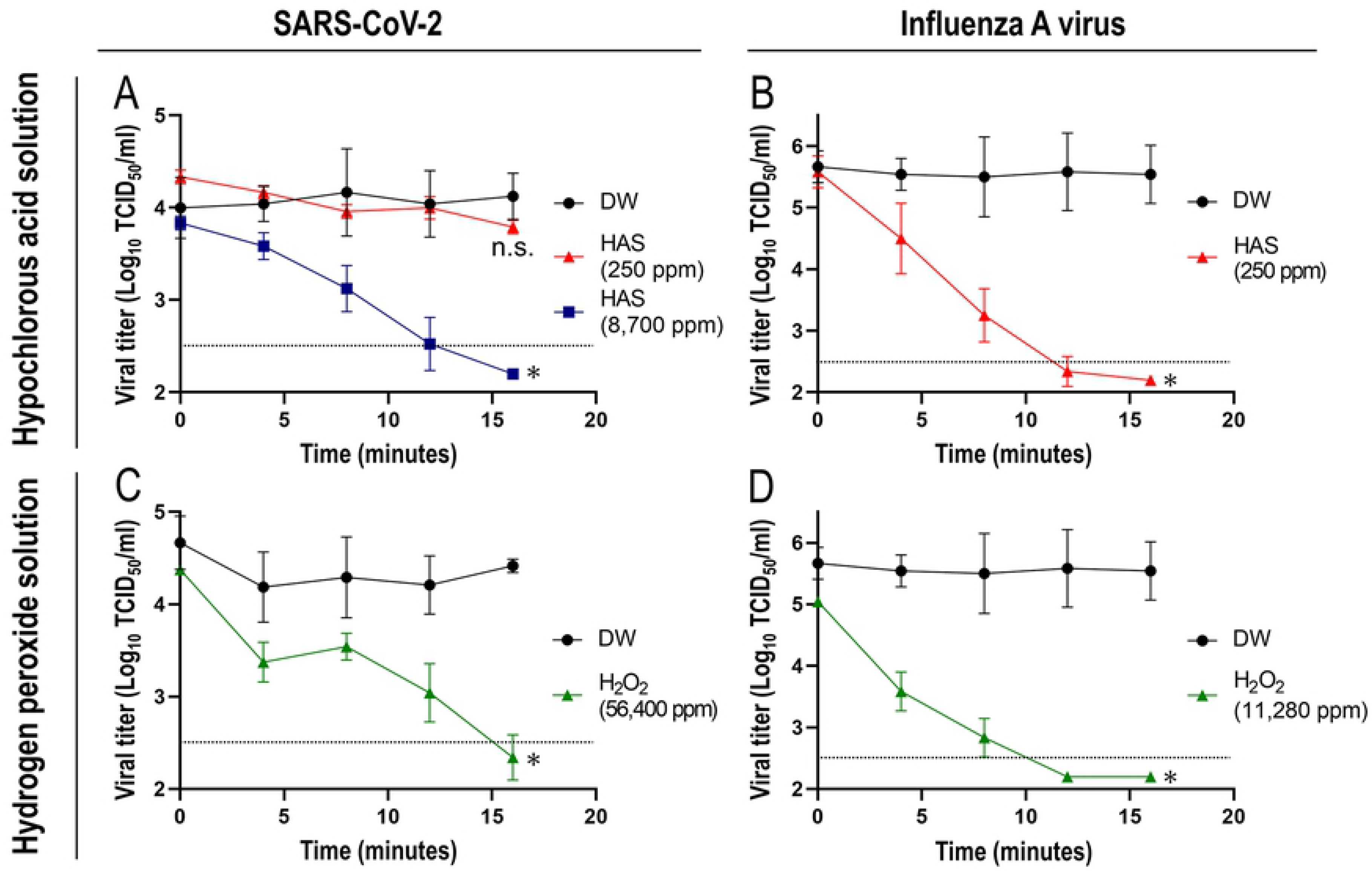
Time-dependent inactivation of viruses by the Dry Fog spraying of disinfectants. SARS-CoV-2 (A and C) and influenza A virus (B and D) were air-dried and exposed to aerosolized hypochlorous acid solution (HAS) (A and B), hydrogen peroxide solution (H_2_O_2_) (C and D), or distilled water (DW) in the form of Dry Fog (A, B, C, and D). The concentrations of spray solutions are as indicated. The viral titer was measured by calculating the TCID_50_ value, as described in the Materials and methods. Horizontal dotted lines in the graphs show the detection limit of the viral titer (3.16 × 10^2^ TCID_50_/ml or 2.5 log_10_ TCID_50_/ml). A viral titer below the detection limit was plotted as half of the detection limit (1.58 × 10^2^ TCID_50_/ml or 2.2 log_10_ TCID_50_/ml). The significance of differences was analyzed using a two-way ANOVA. P < 0.05 was considered to be significant. *P < 0.0001 was estimated for comparisons between DW and HAS 8,700 ppm (A), between DW and HAS 250 ppm (B), between DW and H_2_O_2_ 56,400 ppm (C), and between DW and H_2_O_2_ 11,280 ppm (D). No significant difference was observed between DW and HAS 250 ppm (P > 0.05) (A). n.s. not significant. Each data point represents the average and standard deviation obtained from at least three independent experiments.

### Relationship between spray concentrations and exposed disinfectant amounts

To calculate the exposed disinfectant amount in viral samples in the wells of 96-well microplates, we measured their dissolved concentrations in 20 ml of distilled water in 90-mm Petri dishes during virus inactivation experiments. We then estimated the exposed disinfectant amount in the wells (diameter of 6.5 mm) of a 96-well microplate using an area conversion. Regarding the Dry Fog spraying of hypochlorous acid solution (125, 250, and 8,700 ppm) and hydrogen peroxide solution (1,410, 2,820, 5,640, 11,280, and 56,400 ppm), the total spraying time versus the exposed disinfectant amount as well as the concentration of sprayed disinfectant versus the exposed disinfectant amount per unit spray time are plotted in Figure 4.

**Fig. 4.**
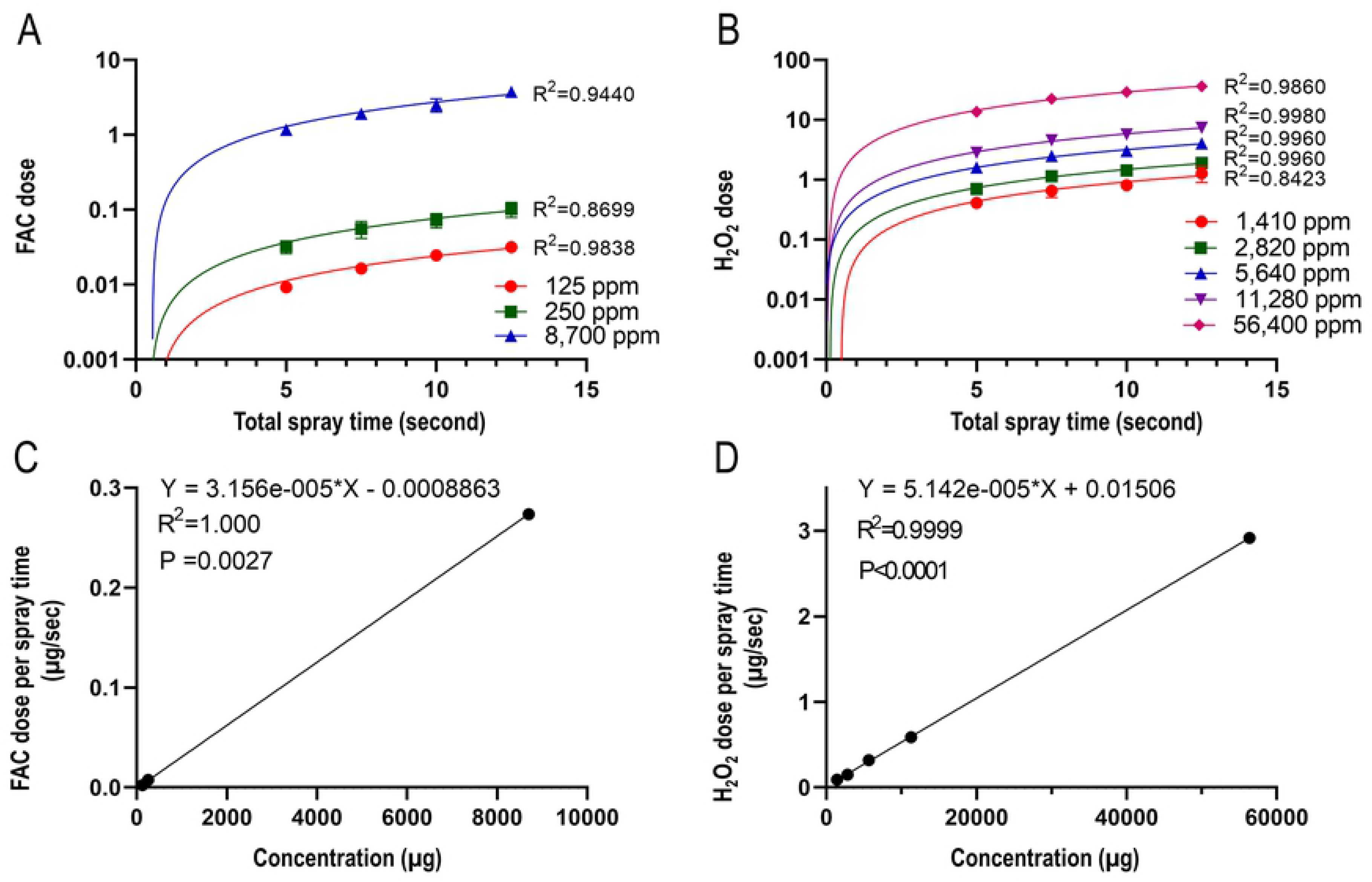
Correlation between sprayed disinfectant concentrations and exposed disinfectant amounts. Hypochlorous acid solution (125, 250, or 8,700 ppm) (A) or hydrogen peroxide solution (1,410, 2,820, 5,640, 11,280, or 56,400 ppm) (B) was sprayed in the form of Dry Fog 4 times 0, 4, 8 and 12 minutes after the initiation of the experiment, as described in the Materials and methods. The exposed disinfectant amounts of free available chlorine (FAC) (A) and hydrogen peroxide (H_2_O_2_) (B) in the wells of a 96-well microplate were calculated and plotted. The exposed disinfectant amounts of FAC (C) and H_2_O_2_ per unit spray time (D) were calculated and plotted. R squared (R_2_) values were estimated using a non-linear regression (curve fit) analysis and reported on the graphs (A and B). In addition, R_2_ values, equations, and p values were estimated using a simple linear regression analysis and reported on the graphs (C and D).

The results obtained confirmed that the spraying of various concentrations of hypochlorous acid solution and hydrogen peroxide solution increased the exposed disinfectant amounts in proportion to the total spraying time (Figs. 4A and 4B). In addition, the exposed disinfectant amount per unit spray time increased in proportion to the sprayed disinfectant concentration of each disinfectant (Figs. 4C and 4D). On one hand, these were expected results because the amount of the disinfectant sprayed from the Dry Fog sprayer was proportional to the total spraying time, and the amount of disinfectant contained in the sprayed droplets was proportional to the concentration. On the other hand, the slope of hypochlorous acid solution was approximately 0.52-fold smaller than that of hydrogen peroxide solution (Figs. 4C and 4D). This result was attributed to the concentration in droplets being lower with the spraying of hypochlorous acid solution [15]. The temperature inside the chamber was maintained at approximately 19-26°C, while humidity was maintained at 59-99% during Dry Fog spraying experiments.

### Relationship between virucidal effects of disinfectants and exposed disinfectant amounts

The relationship between the virucidal effects of aerosolized disinfectants for SARS-CoV-2 and influenza A virus versus the exposed disinfectant amounts of FAC and hydrogen peroxide are shown in Figure 5. The absolute value of the slope of the graph is the virus inactivation constant. With the spraying of hypochlorous acid solution, the inactivation constant of SARS-CoV-2 was approximately 58-fold smaller than that of influenza A virus. In addition, with the spraying of hydrogen peroxide solution, the inactivation constant of SARS-CoV-2 was approximately 9.6-fold smaller than that of influenza A virus. Therefore, SARS-CoV-2 was more resistant to hypochlorous acid solution and hydrogen peroxide solution than influenza A virus.

**Fig. 5.**
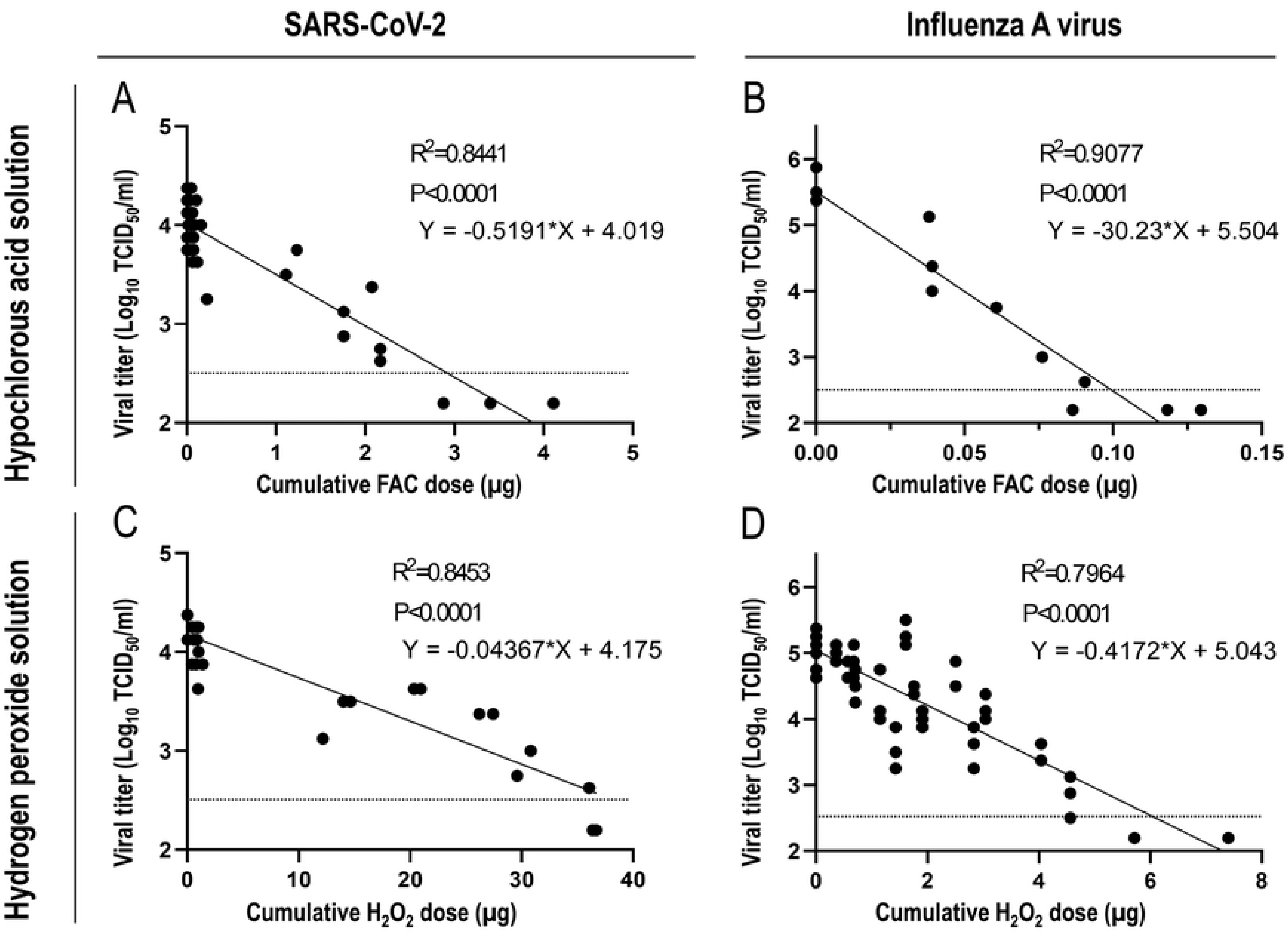
The relationship between the exposed disinfectant amount and virucidal effects of a disinfectant. SARS-CoV-2 (A and C) and influenza A virus (B and D) were air-dried and exposed to aerosolized hypochlorous acid solution (A and B), hydrogen peroxide solution (C and D), or distilled water in the form of Dry Fog (A, B, C, and D). The viral titer was then measured by calculating the TCID_50_ value, as described in the Materials and methods. In addition, the exposed disinfectant amounts of free available chlorine (FAC) (A and B) and hydrogen peroxide (H_2_O_2_) (C and D) in the wells of a 96-well microplate were calculated. A scatter plot was created from each dataset of the TCID_50_ value and exposed disinfectant amount for the combination of the virus and disinfectant. R squared (R_2_) values, equations, and p values were estimated using a simple linear regression analysis and reported on the graphs. Horizontal dotted lines in the graphs show the detection limit of the viral titer (3.16 × 10^2^ TCID_50_/ml or 2.5 log_10_ TCID_50_/ml). A viral titer below the detection limit was plotted as half of the detection limit (1.58 × 10^2^ TCID_50_/ml or 2.2 log_10_ TCID_50_/ml).

### Effects of saliva on the inactivation of SARS-CoV-2 and influenza A virus by the Dry Fog spraying of hypochlorous acid solution and hydrogen peroxide solution

As shown in Figures 3 and 5, the Dry Fog spraying of hypochlorous acid solution and hydrogen peroxide solution inactivated SARS-CoV-2 and influenza A virus. We prepared air-dried viral samples using viral supernatants for those experiments, and viral solutions differed from physiological conditions. Therefore, to assess the effects of saliva components on the virucidal effects of the Dry Fog spraying of disinfectants, dried viral samples were prepared by mixing viral supernatants and artificial saliva solution or PBS as the negative control. Hypochlorous acid solution and hydrogen peroxide solution were then Dry Fog sprayed for viral samples. The results obtained revealed no significant differences in the time kinetics of viral inactivation upon the Dry Fog spraying of disinfectants between dried viral samples prepared with artificial saliva and PBS (Fig. 6).

**Fig. 6.**
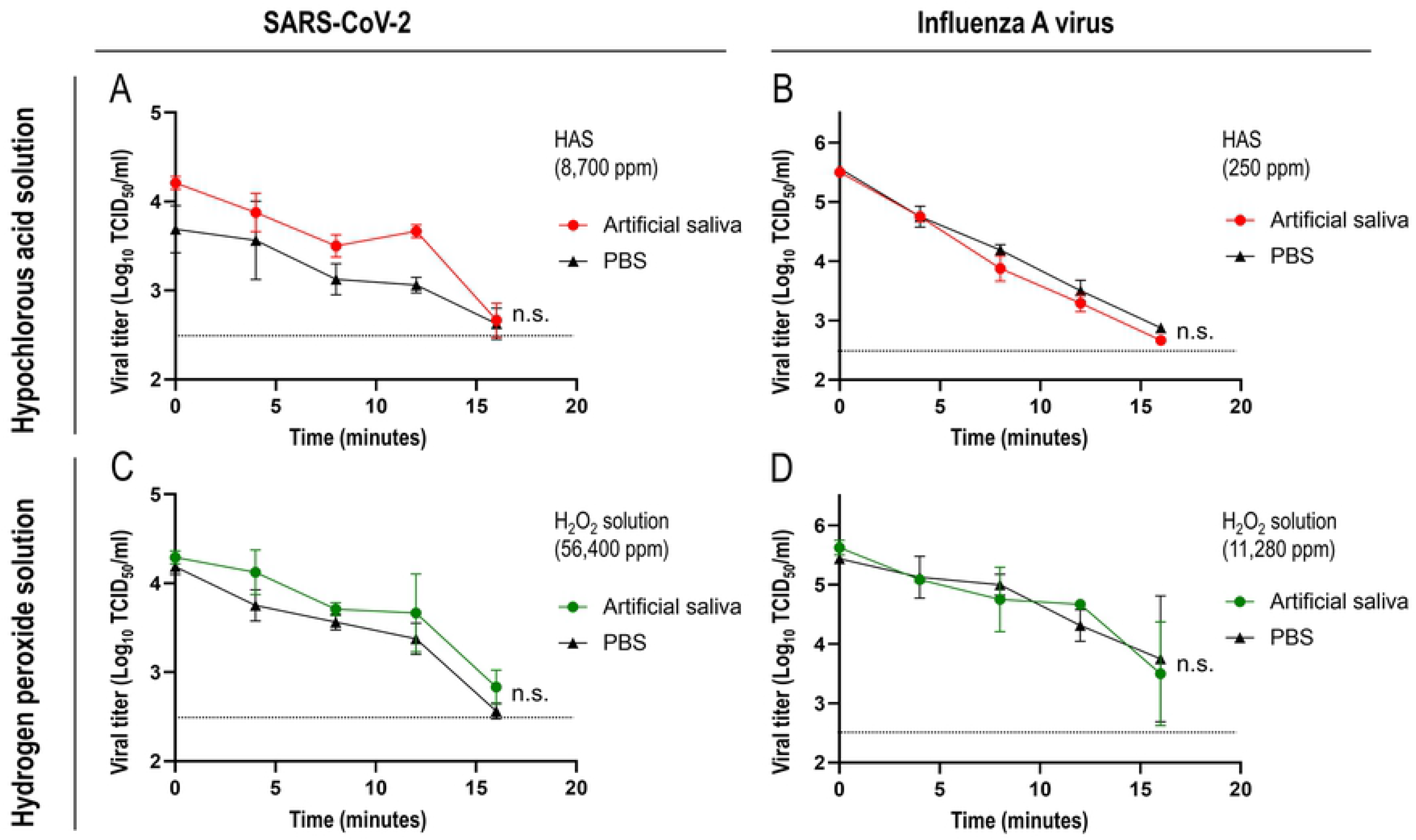
Effects of saliva on the inactivation of SARS-CoV-2 and influenza A virus by the Dry Fog spraying of disinfectants. SARS-CoV-2 (A and C) or influenza A virus (B and D) was mixed with artificial saliva or PBS, air-dried, and exposed to aerosolized hypochlorous acid solution (HAS) (A and B) or hydrogen peroxide solution in the form of Dry Fog (C and D). The disinfectant concentrations of spray solutions are as indicated. The viral titer was measured by calculating the TCID_50_ value, as described in the Materials and methods. Horizontal dotted lines in the graphs show the detection limit of the viral titer (3.16 × 10^2^ TCID_50_/ml or 2.5 log10 TCID_50_/ml). The significance of differences between artificial saliva- and PBS-containing samples was analyzed using an unpaired *t*-test. No significant differences were observed between the samples (P.>0.05). n.s. not significant. Each data point represents the average and standard deviation obtained from at least two independent experiments.

## Discussion

We herein investigated whether SARS-CoV-2 and influenza A virus that had been air-dried and adhered to an environmental surface were inactivated by the Dry Fog spraying of disinfectants. The inactivation of pathogenic viruses, including SARS-CoV-2, influenza A virus, norovirus, and adenovirus, by the Dry Fog spraying of peracetic acid (PAA) or a mixture of PAA and hydrogen peroxide was previously demonstrated [16–21]. Therefore, the Dry fog spraying of disinfectants is considered to effectively inactivate pathogenic viruses on environmental surfaces in laboratories, safety cabinets, and health care facilities. Hypochlorous acid solution and hydrogen peroxide solution, which are unlikely to remain in the environment after spraying, were employed as spray disinfectants in the present study. The results obtained revealed that even though the concentration of the spray disinfectant required for virus inactivation by Dry Fog spraying differed, the infectivities of SARS-CoV-2 and influenza A virus were both reduced to below the detection limit over time with the spraying of disinfectants (Fig. 3). Previous studies reported the high stability of SARS-CoV-2 on environmental surfaces [4–7]. Consistent with these findings, spraying with higher concentrations of hypochlorous acid solution and hydrogen peroxide solution was required to inactivate SARS-CoV-2 than influenza A virus. SARS-CoV-2 and influenza A viruses are both enveloped RNA viruses; however, experiments to elucidate differences in their resistance to the Dry Fog spraying of these disinfectants were not conducted in the present study.

The merits of Dry Fog spraying are 1) it does not wet the environmental surface and 2) it is diffusible and has a small droplet size; therefore, an object will not be excessively wet unless spraying is performed in the same place for a long time. With Dry Fog spraying, it is not necessary to wipe off the disinfectant after spraying, and it is also possible to use it in an environment that cannot become wet. Regarding diffusivity, since the droplet size is small, droplets are assumed to fall at a low speed and float in the air for a long time. By utilizing this property of Dry Fog, it is also possible to diffuse droplets using a fan. Furthermore, droplets behave like air and evenly reach the backs of objects, such as desks and chairs, as well as gaps that cannot be reached; therefore, Dry Fog spraying is considered to be suitable for spatial spraying. On the other hand, one of the disadvantages of Dry Fog spraying is the risk of the inhalation of droplets. Therefore, several factors need to be considered, such as the concentration of droplets in space, the staying time, the amount of suction expressed by the number of breaths, the concentration of the solution in droplets, and the toxicity of the spray solution to the human body. In the present study, we assumed an unmanned space and, thus, did not consider risks to the human body; however, when spraying a disinfectant, such as Dry Fog, in a manned environment, it is extremely important to consider risks to the human body.

The results of the Dry Fog spraying of hypochlorous acid solution and hydrogen peroxide solution using the test chamber confirmed that the exposed disinfectant amount of FAC and hydrogen peroxide per unit spray time increased according to the spray amount and sprayed disinfectant concentration (Fig. 4). Furthermore, SARS-CoV-2 and influenza A virus were inactivated in a manner that was dependent on the exposed amounts of disinfectants (Fig. 5). The present results were obtained from a spraying experiment conducted in a small space (approximately 0.11 m^3^), and, thus, various factors need to be considered when Dry Fog spraying in actual spaces, such as the vaporization of sprayed droplets and a reduction in the disinfectant concentration in droplets due to the longer distance travelled by droplets [22]. In the present study, a maximum of approximately 8 ml of disinfectant was sprayed in a space of approximately 0.11 m^3^, and it is inadequate to simply calculate the amount of disinfectant needed from the volume of the space to be sprayed. It may be necessary to spray more or higher concentrations of a disinfectant in actual spaces. Since the above factors of spraying in actual spaces reflects the exposed disinfectant amount, it is important to measure the exposed disinfectant amount at the site of use in order to confirm its effectiveness.

Countermeasures against infection sources, transmission routes, and susceptible hosts, which are regarded as three important factors for the occurrence of infectious diseases, are considered to be general countermeasures against infectious diseases. WHO recommends avoiding 3Cs (spaces that are closed, crowded or involve close contact), the wearing of a mask, and the frequent cleaning of hands as a measure to block the transmission routes of COVID-19. In addition, vaccination is recommended worldwide as a measure for susceptible hosts to acquire immune protection against the virus. As a countermeasure against infection sources, patient isolation and quarantine are currently being performed. Regarding more active countermeasures against infection sources, the inactivation of SARS-CoV-2 by UV or ozone irradiation has been examined at basic research levels as part of an attempt to inactivate the virus adhering to environmental surfaces [23–25]. The inactivation of influenza A virus adhering to environmental surfaces using an ultrasonic atomizer of hypochlorous acid solution has been reported [26]. In the present study, we examined whether the Dry Fog spraying of hypochlorous acid solution and hydrogen peroxide solution effectively inactivated SARS-CoV-2 and influenza A virus. The results obtained showed that spatial spraying was effective against viruses adhered to plastic microplates, suggesting that spatial spraying is an active countermeasure against infection sources on environmental surfaces.

There are a number of limitations that need to be addressed. Only the SARS-CoV-2 Wuhan strain and influenza A virus H1N1 strain were tested in the present study. SARS-CoV-2 variants of concern, including the delta variant, are continually emerging. Furthermore, there is a wide variety of influenza A viral strains. Inactivation experiments were not performed on these viral strains in the present study. However, a previous study suggested that hypochlorous acid attacks multiple components of microorganisms, including the plasma membrane and nucleic acids, as its germicidal effect [10]. Although the structures of viral proteins due to genetic mutations may differ between the Wuhan strain and other SARS-CoV-2 variants, there may be a commonality in the basic structures and components of virions, such as the lipid bilayer (envelope) and structural and non-structural proteins. Therefore, Dry Fog spraying is expected to act effectively not only against the Wuhan strain and H1N1 strain tested in the present study, but also against other virus strains. As another limitation, we only examined the effects of Dry Fog spraying on viruses dried on the surfaces of plastic microplates. Further studies are warranted to investigate its effects on viruses adhering to the surfaces of other materials. Furthermore, the mechanisms by which Dry Fog spraying affects the infectivities of viruses in a space need to be elucidated. However, difficulties may be associated with establishing an experimental method that safely and quantitatively assesses virucidal effects by recovering the virus in the space after exposing it to Dry Fog, and this has not yet been implemented. Nevertheless, we consider spatial spraying to be an effective method for inactivating SARS-CoV-2 and influenza A virus on environmental surfaces. The accumulation of more information in the future and the development of methods that inactivate pathogens, such as viruses, in the environment for practical use are desired.

## Acknowledgments

We thank Kenji Ayagi, Issei Toyoda, and Masataka Ishida of the Innovation Commercialization Division, Kobe University for their efforts to coordinate the industry-academia joint research project. The manuscript was proofread by Medical English Service, Kyoto, Japan.

## Author contributions

M.U., A.K., and M.K. conceived the study. M.U., T.K., and M.K. performed the experiments. M.U. and M.K took the lead in writing the manuscript. A.K., K.S., and Y.M. provided feedback and helped shape the research and manuscript.

## Notes

### Competing Interest Statement

The authors have declared no competing interest.

